# EthoTrace: from behaviour to neural dynamics in a unified analysis system

**DOI:** 10.64898/2026.10.04.756578

**Authors:** Henry Oo, Can Tao, Guang-Wei Zhang

## Abstract

Understanding how neural populations support natural behaviour requires knowing what a population does while an animal performs a specific, self-initiated action. For responses evoked by an experimenter-delivered stimulus, the signal can be aligned by the trigger itself, and no behavioural scoring is needed to define the event. Complex behaviours such as parental care and social interaction provide no such trigger. They are defined from the video record, usually offline and post hoc, and are conventionally scored in one program and exported to another for analysis. In this study we developed EthoTrace, a single self-contained HTML file that runs in a desktop browser without installation, server or network access and carries a session from behaviour and calcium signal through to the statistical comparison. Behaviour is annotated on a timeline shared with the ΔF/F trace, the signal is corrected and scaled under a stated baseline convention, single bouts stay inspectable in their behavioural context, and repeated bouts are aligned without filling absent data. We demonstrate the complete workflow on a parenting behaviour experiment, in which behaviour was scored and the photometry signal analysed in a single session folder. We expect EthoTrace to facilitate integrative analysis in neuroscience, where behavioural observation and fiber photometry signals are interpreted together.

## Introduction

Fiber photometry resolves population activity from genetically defined neurons in freely moving animals [1]. It is widely used across neuroscience, including studies of social, motivated and parental behaviour [2–5]. Relating its signal to what an animal is doing is how the technique reveals the neuronal substrates of a behaviour. When activity is time-locked to an experimenter-delivered stimulus, the alignment is supplied by the trigger itself. Stimulus timestamps are recorded alongside the signal, and no behavioural annotation is required to define the event. Studies of sensory-to-limbic transformation in the medial septum have used this design, averaging responses directly on the onset of a delivered cue [6,7].

Many behaviours of interest are more complex than a delivered stimulus. Self-initiated social and parental behaviours offer no trigger to align on. Their boundaries exist only in the video record and are often defined post hoc and offline by a human behavioural annotator, as in work on medial preoptic control of parental behaviour and affective state and on prosocial responses to unconscious conspecifics [8–10]. Aligning those annotated behaviours with the simultaneously recorded signal can reveal critical associations between neural activity and specific actions.

Many tools have been developed for each half of this work. Event-logging software supports structured ethograms and keyboard coding [18]. Pose-estimation and supervised or unsupervised classifiers label behaviour automatically at scale [19–30]. Photometry packages provide control-channel correction, event-aligned averaging and transient detection through graphical interfaces [11–16]. They remain separate programs, so the correspondence between behaviour and signal survives only as an exported file [11–17]. Each program carries its own notion of time, so frame rates and event definitions are reconciled by hand. The statistical step is typically written afresh for each project, and the choices it embodies are not recorded with the result. Analytical decisions taken after seeing the data and left unreported are a recognized source of irreproducibility [31,32], and the reporting of computational provenance is correspondingly emphasized [33,34].

In this study we introduce EthoTrace, which integrates behavioural annotation with fiber photometry signal processing in one program in order to support that understanding. EthoTrace is a single HTML file that opens in a desktop browser and reads and writes the session folder in place. It holds behavioural annotation, ΔF/F computation, event alignment, visualization and statistical testing in one environment. Because every operation reads the same annotation, the frame grid, the event definition and the event-to-signal mapping remain explicit components of the analysis rather than assumptions inherited from a file written elsewhere. Incomplete aligned windows are left unfilled and marked rather than padded. For each comparison the tool reports which test it selected and what the test it did not select would have given.

## Results

### EthoTrace is self-contained, from behaviour and calcium signal to analysis

EthoTrace holds behavioural annotation and fiber-photometry analysis in one place. The entire application is a single self-contained HTML file written in plain JavaScript. Opening that file in a desktop browser launches the complete environment. There is no package to install, no Python or MATLAB runtime to maintain, no server component and no account. The file reads and writes the session folder through the browser’s File System Access API, which means the recording is accessed in place on the user’s own computer and no data are uploaded. Annotations and analysis settings are written back into that folder as plain files, so a session carries its own analysis configuration alongside its raw data.

A session folder containing the behavioural video and the photometry file is opened as a unit, and both streams are loaded onto a single timeline rather than being handled by separate programs ( **Fig. 1A–D**). The photometry file supplies a timestamp in milliseconds with paired 410-nm and 470-nm samples on each row. Behaviour is scored on a frame grid whose rate the user sets, independently of the video’s native frame rate. A recording filmed at 60 frames/s can therefore be scored on the 30 frames/s grid used here, without resampling the video. The two streams are represented on a common timebase that starts at zero for both. The analyses place photometry samples by their own timestamps rather than by frame index. In the demonstration session each sample fell within 40 ms of its timestamp. Because the frame grid, the event definition and the event-to-signal mapping are specified within the same environment, they remain explicit components of the analysis rather than assumptions introduced during an intermediate export and re-import step.

**Fig. 1.**
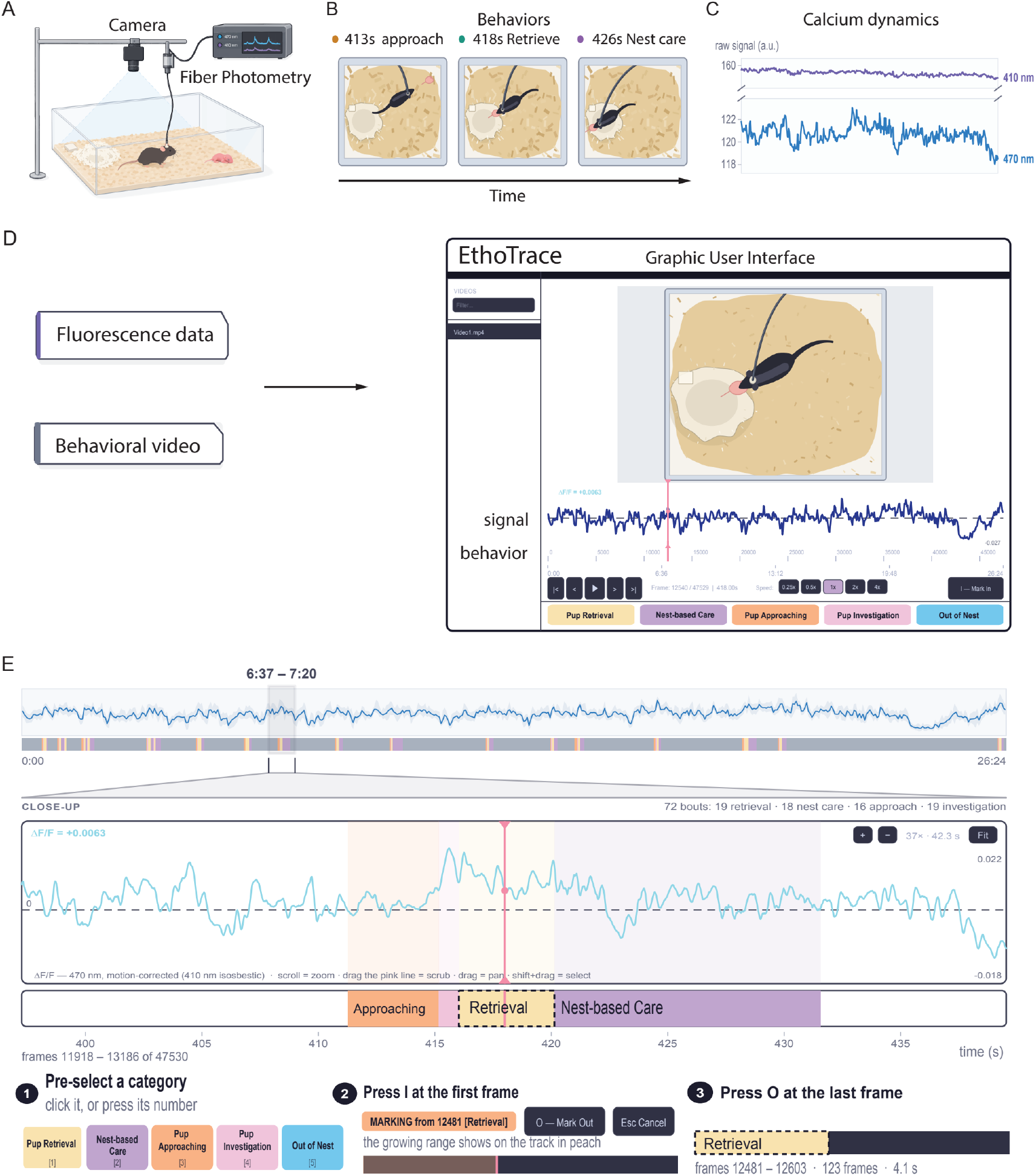
Integrated behavioural annotation and photometry analysis. **A**, Recording setup: a Vgat-Cre mouse with a unilateral fibre implant over MPOA, connected to a fiber photometry system (470-nm excitation, 410-nm isosbestic) and filmed from above (illustration). **B, C**, The two streams acquired simultaneously. **B**, the overhead video (`Video1.mp4`, 1280 × 720, 60 frames/s), shown as three drawn stills at 413, 418 and 426 s. **C**, the fluorescence record (`Fluorescence.csv`), with raw 410- and 470-nm signals over the same interval. **D**, Both streams opened together in EthoTrace (redrawn Annotate tab; video frame drawn), with the display ΔF/F of the whole session, the behaviour timeline and the five scoring categories; playhead at frame 12540 of 47,530 (418.00 s). **E**, Annotation on the shared timeline. Top, the whole 26.4-min recording with all 72 bouts (figure device); middle, close-up of 6:37–7:20 (frames 11918–13186) with the annotation track; bottom, the three steps labelling retrieval Trial 10: select a category, press **I** at the first frame, press **O** at the last. This writes frames 12481–12603 (123 frames, 4.1 s inclusive; 4.07 s in analyses). ΔF/F in **D** and **E** is the display ΔF/F (*F*_0_ = fitted 410-nm signal), ≈0.010 below the analysis ΔF/F in Figs 2–4.

### EthoTrace annotates both streams on a synchronized timeline

Behaviour is scored within that shared timeline ( **Fig. 1D, E** ). The video plays above the ΔF/F trace and the behaviour track, all three sharing one zoom window, so the experimenter navigates signal and behaviour together. Here we demonstrate the system on parenting behaviour, so the categories used throughout are the ethogram for that assay: five categories assigned to keys 1–5, namely pup retrieval, nest-based care, pup approaching, pup investigation and out of nest, four of which were used in the demonstration dataset. The category list is user-definable: a study of other behaviours defines its own ethogram in place of this one, and the tool imposes no restriction on which behaviours are scored. A bout is defined by pressing **I** (in) at its first frame and **O** (out) at its last. The category can be selected before or after marking. Bouts occupy one exclusive track, so no frame can carry two labels; they can be relabelled, resized or deleted with undo, and each can be exported as a video clip for inspection or for training an automated classifier elsewhere. Annotations save automatically as a JSON file listing each bout’s inclusive frame range and category. The ΔF/F trace can be hidden, which is how the demonstration session was scored so that labelling was blind to the signal.

The same annotation drives every subsequent operation: ΔF/F computation, event alignment, visualization and statistical testing. Relating behaviour to the photometry signal therefore requires no separate linking step. That linking step is ordinarily where an annotation program hands off to an analysis script, and it is where frame-rate and alignment errors enter without announcing themselves.

A pup-retrieval session recorded in a Vgat-Cre dam with a unilateral fibre implant over the medial preoptic area exercises the system end to end. Across the 26.4-min session, 72 bouts of approach, investigation, retrieval and nest-based care were scored on the 30 frames/s grid covering 47,530 frames ( **Fig. 1E**). The dataset serves to demonstrate each stage of the workflow rather than to support a biological claim.

### EthoTrace computes ΔF/F with a stated baseline convention

Within the same environment, the 470-nm signal is corrected for calcium-independent fluctuations using the 410-nm isosbestic channel. The signal is regressed on the isosbestic channel by ordinary least squares across the whole session. The baseline is the fitted control, offset by the 10th percentile of the regression residuals. A Savitzky–Golay filter smooths the resulting ratio. In the demonstration session the isosbestic fit gave *r* = 0.95.

Two consequences of this convention are worth reporting, because both are ordinarily left implicit. First, the percentile offset is what makes ΔF/F read positive. Least-squares residuals average zero. Taking the fitted control itself as the baseline would therefore centre ΔF/F on zero, leaving 47.8% of samples negative in this session. The offset instead sets ΔF/F = 0 at the 10th percentile of the residuals. 90.3% of analysis ΔF/F samples were then positive. Here that offset was −0.97% of the fitted control. Second, the smoothing is mild but not negligible. At the 30.013-Hz per-channel sampling rate the 11-sample window spans 0.37 s, with a −3-dB point of 3.0 Hz. It reduced the standard deviation of sample-to-sample differences by 60%. It lowered the peak of the mean retrieval-aligned trace by only 0.5%. Removing either the filter or the percentile offset changed neither the tests selected nor the significance marks reported below, so the statistical conclusions in this session do not rest on either choice. No other detrending, *z*-scoring or per-trial baseline subtraction is applied.

One distinction follows from the need for a responsive interface. The Annotate tab displays a *display ΔF/F* computed against the fitted control alone, which in this session lies approximately 0.010 below the *analysis ΔF/ F* used for all statistics. The display trace serves navigation and annotation; the analysis trace serves every reported number. Keeping the two explicitly separate avoids a class of error in which a trace read off the annotation screen is later treated as the analysed signal.

### EthoTrace keeps single trials inspectable in behavioural context

An annotated bout becomes an analysis object as soon as it is marked, so individual events can be examined before any averaging (**Fig. 2**). Every retrieval bout is drawn as a trial, numbered in annotation order, within one fixed window from 5 s before to 10 s after its onset, and all annotated bouts falling in that window are shaded by behaviour in place. The retrieval marked in **Fig. 1E** appears unchanged as Trial 10. Its onset sits at *t* = 0 and its observed offset at 4.07 s. The preceding approach and investigation, and the subsequent nest-based care, remain drawn around the same photometry trace ( **Fig. 2B, C** ). The whole-session view above it retains the same information at session scale, with every retrieval window shaded against the continuous trace and all 72 bouts in a ribbon beneath.

**Fig. 2.**
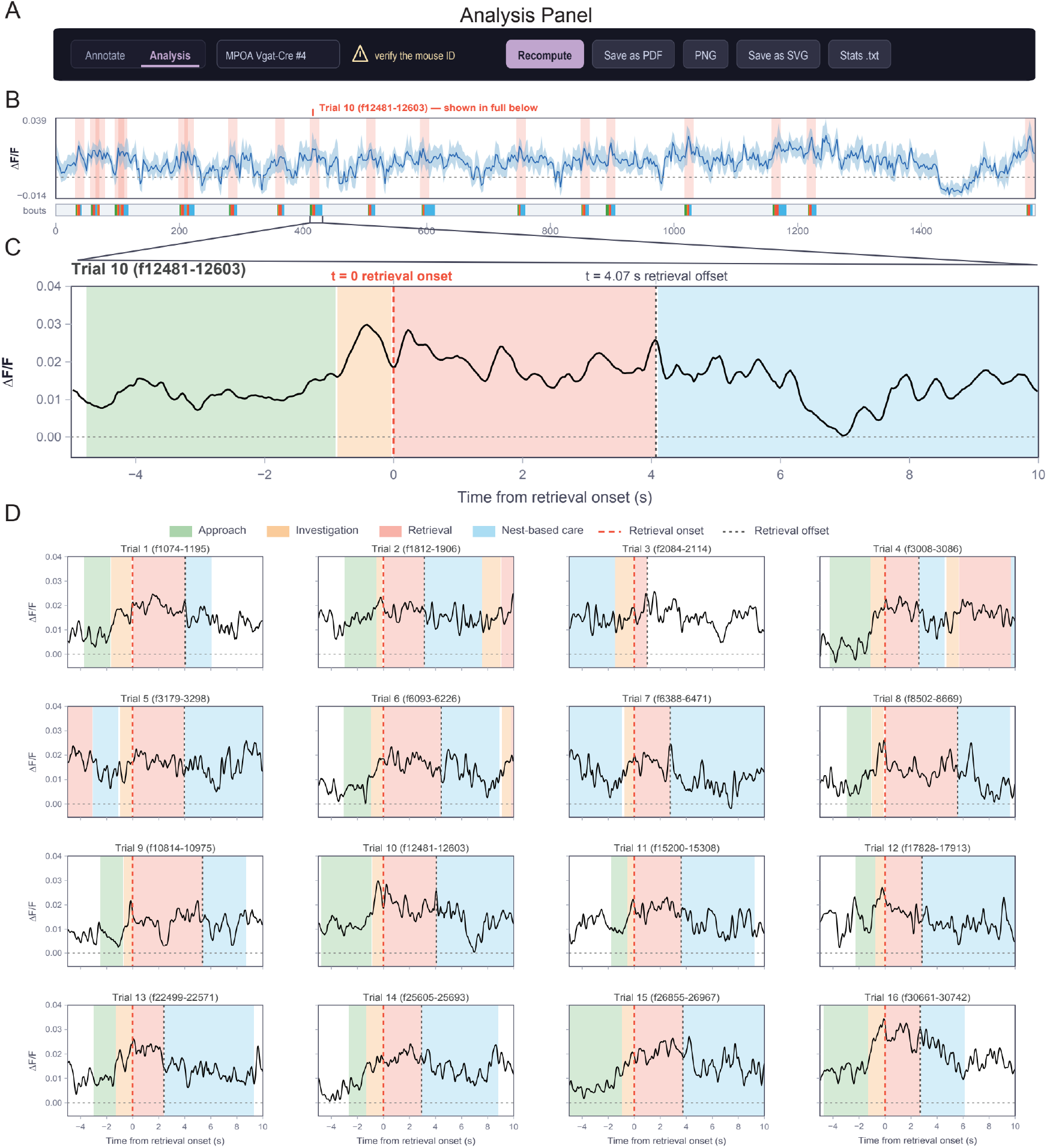
Event-aligned photometry during individual pup retrievals. **A**, Analysis toolbar with this session’s settings (automatic test selection; look-back, 16.5 s), simplified and redrawn; explanatory text added for the figure, and the panel letters it names refer to EthoTrace’s own report. Neither setting affects **B**–**D. B**, Analysis ΔF/F across the 26.4-min session (line, bin mean; band, bin minimum–maximum; dashed line, zero), with the −5 to +10 s window around each retrieval onset shaded and all 72 bouts in the ribbon below; Trial 10, annotated in Fig. 1E, is marked. **C**, Trial 10 (frames 12481–12603) at single-bout resolution, with retrieval onset at *t* = 0 (dashed line), observed offset at 4.07 s (dotted line), and surrounding approach, investigation and nest-based care shaded. **D**, The first 16 of the session’s 19 retrieval trials, in annotation order on a shared *y*-axis, redrawn from report panel E; the remaining three are drawn in EthoTrace’s own report and are omitted here for space.

Behaviours in pup retrieval occur as a stereotyped sequence. A signal change near an event onset may belong to the scored behaviour or to what preceded it. An average across trials can obscure whether a signal change is associated with the scored behaviour or with the one that preceded it. Drawing each trial with its own local context makes that ambiguity visible on the trial itself ( **Fig. 2D** shows the first 16 of the 19 retrievals). A single event can then be read as an event, not as a segment of trace.

*EthoTrace aligns bouts without filling absent data*

The system then aligns repeated bouts of any annotated behaviour and summarizes them without discarding the observations behind the summary ( **Fig. 3** ). Bouts of a behaviour are aligned to their own onsets. Each is interpolated onto a grid at the photometry sampling interval, from a user-set pad before onset to a pad after *each bout’s own offset*. A bout’s full observed duration is therefore represented, not truncated to a common length. The bouts are drawn as a heat map, sorted longest first, on one symmetric colour scale shared across all four behaviours. The mean ± SEM beneath is computed at each time point over the bouts that have data there ( **Fig. 3B, C**).

**Fig. 3.**
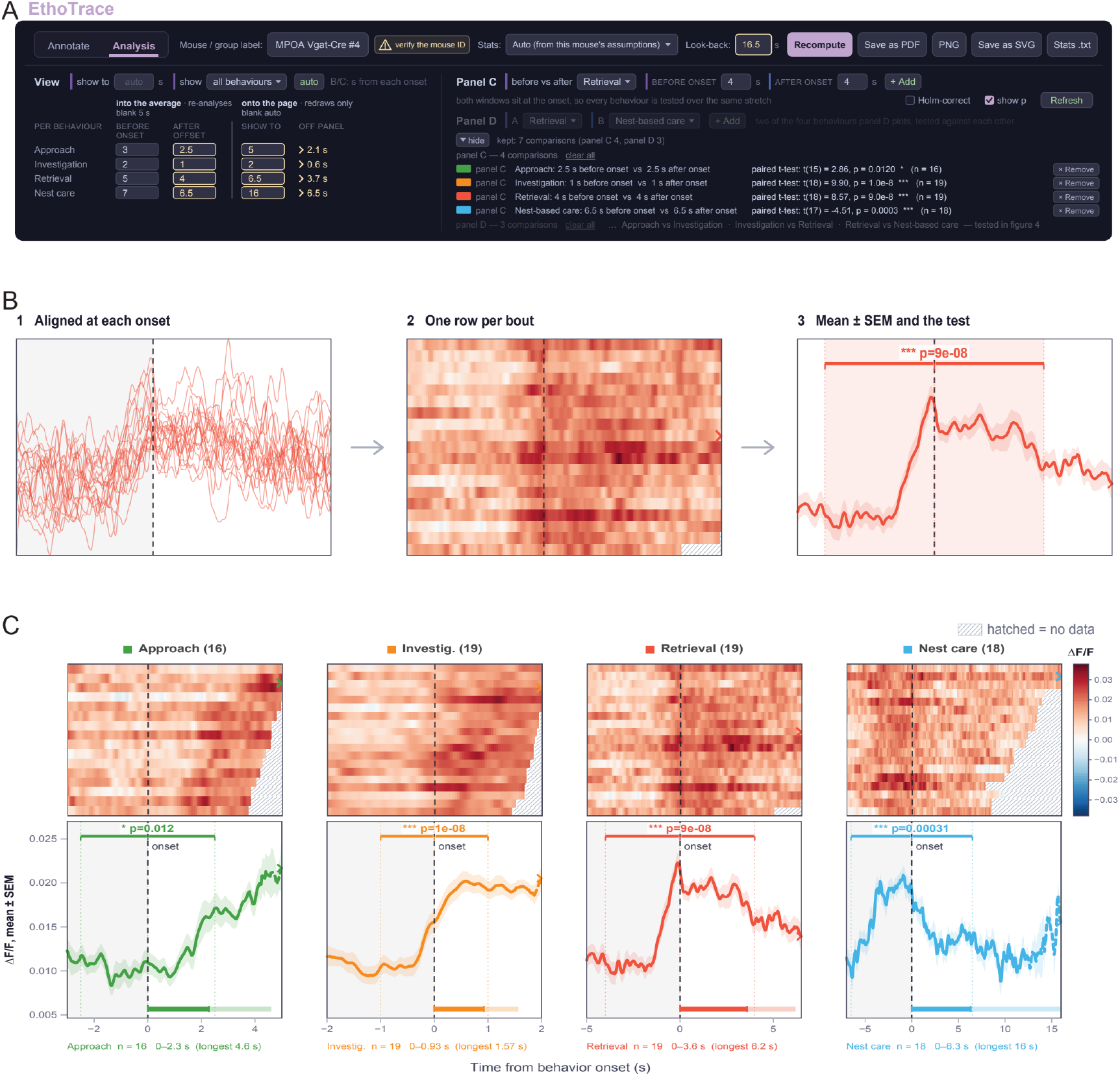
Behaviour-aligned photometry across annotated bouts. **A**, Analysis tab with this session’s settings, redrawn: per-behaviour pads before onset and after offset (3/2.5, 2/1, 5/4 and 7/6.5 s), display limits (“show to”: 5, 2, 6.5 and 16 s) with the extent of data beyond them (“off panel”), the four before/after comparison windows (2.5, 1, 4 and 6.5 s), and Holm correction off; dimmed controls belong to Fig. 4. **B**, Construction of an aligned summary: bouts aligned at each onset, one heat-map row per bout, then the mean ± SEM with the compared windows tinted. **C**, Heat maps (rows sorted longest first; hatched cells, no data) above the mean ± SEM (dashed where fewer than half the bouts contribute) for approach ( *n* = 16), investigation (Investig., *n* = 19), retrieval (*n* = 19) and nest-based care (Nest care, *n* = 18), each on its own time axis with one shared ΔF/F colour scale and *y*-axis. Brackets span the compared windows (dotted lines) with uncorrected paired *t*-tests (*t*(15) = 2.86, *P* = 0.012; *t*(18) = 9.90, *P* = 1.0 × 10^−8^; *t*(18) = 8.57, *P* = 9.0 × 10^−8^; *t*(17) = −4.51, *P* = 3.1 × 10^−4^); bars run from onset to the mean (solid) and longest (pale) bout duration. Grey shading, pre-onset pad; chevrons, data beyond the display limit. * *P* < 0.05; *** *P* < 0.001. These statistics demonstrate the software workflow within one session and are not population-level biological inference.

Because bouts differ in length, most aligned windows extend beyond the data available for the shorter bouts, and this is the common case rather than an edge case. Those regions are left empty rather than padded, held at the last value or extrapolated: they appear as hatched cells in the heat map, and the population mean is drawn dashed wherever fewer than half the bouts contribute. Bout durations differ by an order of magnitude across the four behaviours in this session. Sorting rows by duration exposes the changing coverage directly as a staircase. A per-behaviour display limit controls how far after onset each column is drawn without altering the average, which lets a 16-s behaviour and a 1-s behaviour be shown legibly on one figure.

Each behaviour was compared before versus after its own onset, over matched windows set per behaviour. Automatic selection chose a paired *t*-test in all four cases. The test statistics are given in the **Fig. 3** legend and are not corrected for multiple comparisons.

### EthoTrace reports the test it selected and the one it did not

Finally, the system reduces each bout to a single measurement and carries the statistical reasoning through to the reported comparison (**Fig. 4**). A bout contributes the mean of its analysis ΔF/F between its observed onset and offset. For Trial 10 that mean was 0.01944, from the 122 samples spanning its 4.07 s ( **Fig. 4B**). The surrounding ±3 s of context is displayed but excluded from the mean. The resulting per-bout values form the distributions compared between behaviours as box plots with every bout shown as an individual point (**Fig. 4C**).

**Fig. 4.**
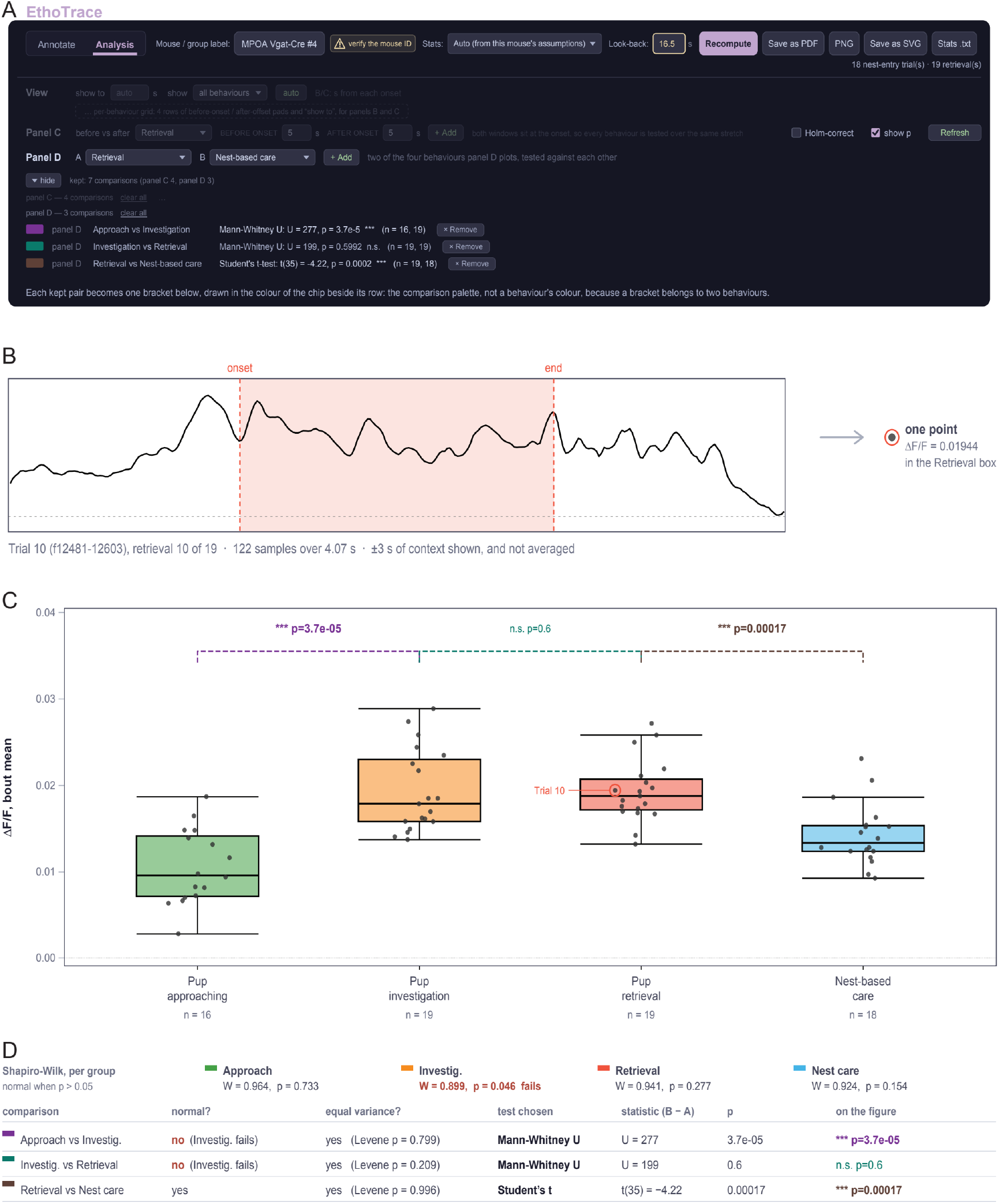
Bout-level photometry measurements and statistical comparisons. **A**, Analysis-tab controls for report panel D, redrawn, listing the three kept pairs with their tests and uncorrected *P* values; dimmed controls belong to Fig. 3, and the explanatory sentence was added for the figure. **B**, One bout, one measurement: the mean of Trial 10’s analysis ΔF/F from onset to offset (0.01944, from 122 samples over 4.07 s); the ±3 s of context is displayed but not averaged, and the dotted line marks zero. **C**, Per-bout mean ΔF/F (*n* = 16, 19, 19 and 18 bouts) as Tukey box plots (whiskers, 1.5 × IQR) with every bout as a jittered point and Trial 10 marked; dashed brackets, coloured as the chips in **A**, show the kept pairs. **D**, Decision path: Shapiro–Wilk normality per behaviour (*W* = 0.964, 0.899, 0.941 and 0.924; *P* = 0.733, 0.046, 0.277 and 0.154) and median-centred Levene tests of equal variance [43] per pair (*P* = 0.799, 0.209 and 0.996), the test each combination selects, and its statistic and *P* value. Investigation fails normality at *P* = 0.046, so both of its pairs use the Mann–Whitney *U* test (*U* = 277, *P* = 3.7 × 10^−5^; *U* = 199, *P* = 0.60), while retrieval versus nest-based care uses Student’s *t*-test (*t*(35) = −4.22, *P* = 1.7 × 10^−4^). Statistics are oriented B − A, the second behaviour of each pair minus the first. \*\*\**P* < 0.001; n.s., not significant. These statistics demonstrate the software workflow within one session and are not population-level biological inference.

The test applied to each pair is not fixed in advance. It is selected from assumption checks computed on those same values. Shapiro–Wilk tests on each group and a median-centred Levene’s test choose between Student’s *t*-test, Welch’s *t*-test and the Mann–Whitney *U* test. Both the checks and the resulting selection are reported with the comparison (**Fig. 4D**). In this session one group of bout means narrowly failed the normality criterion at *P* = 0.046 while the other three passed. That single outcome selected a nonparametric test for both comparisons involving that behaviour, and a parametric test for the third. Variances were equal in all three pairs, so normality alone determined each selection. The test statistics are given in the **Fig. 4** legend.

Three features of that selection are recorded. When an assumption test that decides the choice returns *P* between 0.04 and 0.06, the choice is flagged as marginal. The *P* value of the test *not* selected is reported alongside it. A decision that turned on the third decimal place is then visible as such. Test selection can instead be forced to be parametric or nonparametric for a whole session, and the forced choice is recorded rather than silently applied. And a plain-text report lists every assumption check, every test and every excluded bout, together with analyses the tool computes but does not draw. Each comparison is reported two-sided at α = 0.05 with an effect size and a 95% confidence interval of the mean difference, and an optional Holm correction can be applied across the comparisons shown on a panel.

A single assumption test close to the conventional threshold therefore set the statistical path for two of three comparisons in this session. EthoTrace reports which test it selected and what the test it did not select would have given.

## Discussion

What a population signal means during self-initiated behaviour depends on where that behaviour is judged to begin and end. A calcium transient near an onset can belong to the scored action or to whatever preceded it, and in behaviours that unfold as stereotyped sequences, such as parental care, those two possibilities can be separated by a second or less. Which of them a study reports is therefore determined by the annotation as much as by the photometry. Experiments that deliver a stimulus avoid the problem, because alignment comes from the trigger and the event needs no scoring [6,7]. Studies of social and parental behaviour have no such trigger [8–10], so the behavioural boundary becomes part of the measurement, and its precision sets what the signal can be said to show. Aligning behaviour and signal is thus not a formatting step before analysis; it is where the association between neural activity and action is actually established.

EthoTrace is built around that point. It performs behavioural annotation, control-channel correction, ΔF/F computation, event alignment, visualization and statistical testing in one program. Behaviour is scored on a timeline shared with the ΔF/F trace, and every later operation reads that same annotation, so the frame grid, the event definition and the event-to-signal mapping stay explicit instead of being inherited from a file written elsewhere. Aligned windows are retained only where data exist, so thin coverage is displayed rather than filled, and each bout becomes one measurement, with the tool reporting which test it selected for a comparison and what the test it did not select would have given. Compared with five published open-source photometry packages and the reference behavioural event-logger, the feature sets differ in function rather than in completeness (**Fig. 5**, scored from each tool’s own publication [11–15,18]). Those packages provide capabilities EthoTrace does not, including transient detection and batch processing across sessions [11–13,15], and the event-logger supports an unlimited project-defined ethogram with multiple subjects and modifiers [18]. The closest existing bridge between the two halves is PhAT, which imports behaviour in the event-logger’s format on the assumption that both files share a zero point [14]. EthoTrace makes the same assumption, and what it removes is the intermediate export and re-import.

**Fig. 5.**
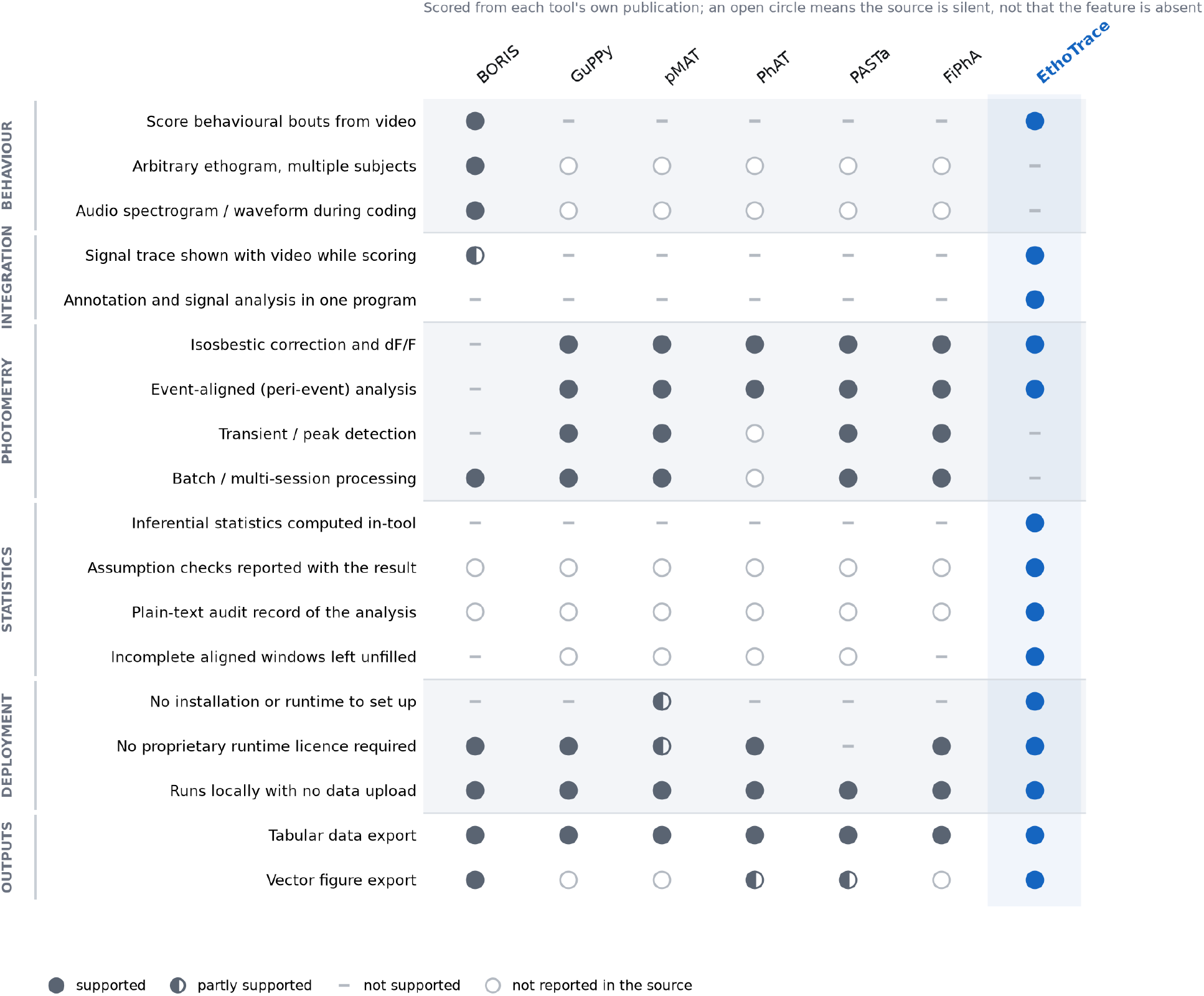
Feature comparison with published open-source tools. Eighteen features scored for the reference behavioural event-logger (BORIS [18]), five open-source photometry packages (GuPPy [11], pMAT [12], PhAT [14], PASTa [13], FiPhA [15]) and EthoTrace. Filled circle, supported; half-filled, partly supported; dash, not supported; open circle, not reported in the source consulted. Scores were taken from each tool’s primary publication, and for BORIS additionally from its user guide, which documents synchronous plotting of up to two external data series. An open circle therefore indicates that the source is silent on that feature, not that the software lacks it; all six tools are actively developed and current versions may exceed their published descriptions. Per-cell evidence and the source for every score are given in Extended Data Table 1. The comparison is a feature survey, not a performance benchmark.

Several properties bound what EthoTrace supports. It analyses one session at a time with the bout as the unit of analysis, so it supports no inference across animals, and manual frame-by-frame scoring does not scale to cohorts, where pose-based and automated classifiers are the appropriate instrument [19–30]. Synchronization assumes both streams start together, which an experiment with TTL alignment would improve on. The isosbestic correction is a convention with known failure modes rather than a guaranteed control: it does not correct the haemoglobin absorption artifact [35], and published guidance stresses checking its adequacy per preparation [17,36]. The percentile offset that makes ΔF/F read positive is a reporting choice, not a physical baseline. The numerical verification is a property of the build described in Methods rather than a warranty on later builds, which is why the reference implementation and the checking procedure are distributed with the tool.

## Methods

### Software architecture

EthoTrace is a single self-contained HTML file written in plain JavaScript, with two tabs, Annotate and Analysis. It reads and writes the session folder through the browser’s File System Access API; annotations and analysis settings ( `analysis_settings.json`) are saved as plain files in that folder, and reports can be exported as PDF, PNG or SVG. The ΔF/F computation and statistical tests reimplement the NumPy [37] and SciPy [38] routines of an in-house Python reference implementation of the same analysis. EthoTrace assumes only a 470-nm signal channel and a 410-nm isosbestic control, and its processing contains no region- or genotype-specific steps. It has been used and tested in Microsoft Edge and Google Chrome, including automated end-to-end tests of development builds in headless mode.

### Input data and synchronization

Each session folder contains the behavioural video (MP4) and the photometry file exported by the acquisition software (`Fluorescence.csv`), in which each row holds a timestamp in milliseconds and paired 410-nm and 470-nm samples. Behaviour is scored on a frame grid whose rate is set by the user (default 30 frames/s), independently of the video’s native frame rate. Bout onsets and offsets are converted to seconds by dividing their start and end frame indices by that rate, so a bout lasts (end − start)/rate; the Annotate tab instead counts a bout’s frames inclusively.

Behaviour and photometry are mapped onto a common timebase that starts at zero for both streams, without offset correction. The analyses place photometry samples by their timestamps, whereas the Annotate-tab trace is spread evenly across the video’s frames, which in the session shown placed each sample within 40 ms of its timestamp. Where a regular time grid is needed, ΔF/F is linearly interpolated between recorded samples, and points not covered by the recording are left empty rather than padded or extrapolated.

### Behavioural annotation

The Annotate tab displays the video above a synchronized ΔF/F trace and a behaviour timeline sharing one zoom window. Categories are user-definable and are assigned in order to keys 1–5. For this study the list was pup retrieval, nest-based care, pup approaching, pup investigation, and out of nest; a study of other behaviours defines its own list in place of these. A bout is labelled by pressing **I** (in) at its first frame and **O** (out) at its last, with the category selected before or after marking. Bouts form one exclusive track, so no frame carries two labels; they can be relabelled, resized or deleted, with undo. Annotations are saved automatically as a JSON file listing each bout’s inclusive frame range and category, and the category list in force is written to the session folder (`category_idx.txt`), so the ethogram used travels with the data. Each bout can be exported as a video clip. The ΔF/F trace can be hidden when scoring should be blind to the signal.

### Photometry processing

The 470-nm signal ( *F*_470_) is corrected for calcium-independent fluctuations with the 410-nm isosbestic signal (*F*_410_) [3,39]. *F*_470_ is regressed on *F*_410_ by ordinary least squares over the whole session, giving the fitted control *F*_pred = β_1_*F*_410_ + β_0_ and the residual *e* = *F*_470_ − *F*_pred. The baseline is *F*_0_ = *F*_pred + *P*_10_(*e*), where *P*_10_(*e*) is the 10th percentile of the residuals, and the analysis ΔF/F is SG[( *F*_470_ − *F*_0_)/*F*_0_], where SG is a Savitzky–Golay filter [40] with an 11-sample window and a cubic polynomial. Both settings were inherited unchanged from the Python reference implementation and were fixed before the session shown was recorded. Two ΔF/F traces exist: for responsive display the Annotate tab shows a display ΔF/F computed with *F*_0_ = *F*_pred, while all statistics use the analysis ΔF/F defined above. No other detrending, *z*-scoring or per-trial baseline subtraction is applied.

### Event-aligned analyses

The Analysis tab produces a three-page report for each session. Four behaviours are analysed: approach, investigation, retrieval and nest-based care. Every retrieval bout is shown as a trial, numbered in annotation order, in a fixed window from 5 s before to 10 s after its onset, with all annotated bouts in that window shaded by behaviour. For behaviour-aligned averages, the bouts of each behaviour are aligned to their own onsets. Each is interpolated onto a grid at the photometry sampling interval, from *T*_pre before onset to *T*_post after each bout’s own offset. The pads are set per behaviour. A per-behaviour display limit sets how far after onset each column is drawn and does not change the average. Bouts are displayed as a heat map, longest first, on one symmetric colour scale shared by all behaviours, and as the mean ± SEM (population SD/√n) at each time point over the *n* bouts with data there. For between-behaviour comparisons, each bout is reduced to the mean of its analysis ΔF/F from onset to offset.

### Statistical analysis

All statistics are computed within one session, with the bout as the unit of analysis. EthoTrace does not compare animals, and because bouts from one recording are not independent, the tests describe that session rather than a population. In the before/after test, each bout contributes its mean ΔF/F over a user-set window before onset and another after onset. Both windows are anchored at onset. Bouts whose aligned traces do not cover both windows are excluded. At least three bouts are required. A Shapiro–Wilk test [41] on the paired differences selects a paired *t*-test (*P* > 0.05) or a Wilcoxon signed-rank test [42]. In the between-behaviour test, the bout means of two behaviours are compared as two groups with a two-sample test: Shapiro–Wilk tests on each group and a median-centred Levene’s test [43] select Student’s *t*-test, Welch’s *t*-test [44] or the Mann–Whitney *U* test [45].

An assumption test returning *P* between 0.04 and 0.06 flags the choice as marginal, and the *P* value of the test not chosen is reported with it. EthoTrace does not imply that assumption-driven branching is an optimal inferential strategy; the implementation shows how such a decision can be made explicit and recoverable where it is used. Selection can instead be forced to be parametric or nonparametric for a session. The forced choice is recorded. A plain-text report ( `Stats.txt`) lists every assumption check, every test and every excluded bout, together with automatic analyses that are not drawn. An optional Holm correction [46] adjusts *P* values across the comparisons shown on a panel. *P* values are not adjusted for the choice of windows or comparisons, nor for assumption-based test selection. Tests are two-sided with α = 0.05, each reported with an effect size (Cohen’s *d*_z, Hedges’ *g* or the rank-biserial correlation) and a *t*-based 95% confidence interval of the mean difference.

### Numerical verification

Because the statistical routines are a dependency-free JavaScript reimplementation rather than calls into an established library, their output was verified against SciPy 1.17.1 and NumPy 2.4.3. Reference values came from the in-house Python implementation of the same analysis, for six other recording sessions, and from SciPy directly for synthetic inputs. The synthetic set included 488 two-sample comparisons constructed with ties and unequal group sizes. *P* values agreed within 2 × 10^−13^ relative error for the *t*, Wilcoxon signed-rank, Levene and Mann–Whitney tests, and within 6 × 10 ^−9^ absolute error for the Shapiro–Wilk test. The session shown was checked the same way end to end: the per-bout values entering each test were passed from the analysis engine to the SciPy reference routines, the seven *P* values agreed within 10^−14^ relative error, the assumption tests selected the same tests, and the analysis ΔF/F agreed within 6 × 10 ^−16^ with the trace computed by the reference implementation. The agreement is at the level of the computation rather than of rounded reported values, with the Shapiro–Wilk statistic the least close of the set. It is a property of this build, not a guarantee for later builds, which is why the reference implementation and the comparison procedure are distributed with the tool.

### Animal procedures

All procedures were approved by the Virginia Commonwealth University Institutional Animal Care and Use Committee and conducted in accordance with the *Guide for the Care and Use of Laboratory Animals* and the Public Health Service Policy on Humane Care and Use of Laboratory Animals. Controlled substances were handled under the responsible investigator’s DEA registration.

A 3.5-month-old female vesicular GABA transporter (Vgat)-ires-Cre knock-in mouse [47] (C57BL/6J background; JAX stock no. 028862) underwent stereotaxic surgery. Anaesthesia was isoflurane (5% induction, 1–2% maintenance; R540, RWD Life Science). Body temperature was maintained (ThermoStar, RWD Life Science). Extended-release buprenorphine (Ethiqa XR, 1.3 mg/mL; Fidelis Animal Health) was given at the start of surgery. An 80-nL volume of pGP-AAV-Syn-FLEX-jGCaMP8s-WPRE [48] (Addgene 162377-AAV1; RRID: Addgene_162377) was delivered to the medial preoptic area of the left hemisphere (AP +466 µm, ML −533 µm, depth 4736 µm). The injection used a pulled glass micropipette (World Precision Instruments capillary; P-97 puller, Sutter Instrument) driven by a nanolitre injector at 20 nL/min. A 6-mm fibre-optic cannula (Inper) was implanted at the same coordinates to a depth of 4536 µm. For histological verification of implant placement, 100-µm sections were labelled with NeuroTrace fluorescent Nissl stain (Thermo Fisher Scientific) and imaged at 2× with multicolour acquisition and stitching (BZ-X1000, KEYENCE).

### Photometry and behavioural recording

Fluorescence was recorded with a camera-based R821 Multichannel Fiber Photometry System (RWD Life Science). It alternated 410-nm and 470-nm excitation at ≈60 Hz, giving 30.013 Hz per channel. Each channel’s signal was the mean intensity of a region of interest over the fibre end face. Both LEDs were set to 30 in software units. Behaviour was filmed from a fixed top-down camera at 1280 × 720 and 60 frames/s (NexiGo). No TTL markers were recorded, so synchronization relies on both streams starting together. The end of the photometry record and the video duration agreed within 7.3 ms. Per-frame camera timing and the frame accuracy of browser video seeking were not independently verified.

### Pup retrieval assay and scoring

Testing took place in the dam’s home cage. Pups (postnatal day 6) were removed before recording. Each presentation began with the dam in the nest, when one pup was placed on the bedding opposite the nest, and ended when the dam released the pup in the nest. Approach began when the dam started walking toward the pup, and ended when she reached it. Investigation began when she reached and briefly sniffed the pup, and ended when she picked it up. Retrieval began when she lifted the pup, and ended when she released it in the nest. Nest-based care began at that release, and ended when she left the nest. In three of 16 presentations the dam released the pup at the nest edge, then returned later to investigate and retrieve it to a new location. These second retrievals were scored as investigation and retrieval without a preceding approach. The session therefore contains 19 retrievals. The experimenter scored the video with the ΔF/F trace hidden. The out-of-nest category was not scored, because the interval between presentations varied.

### Dataset and figure preparation

All data come from one recording session of 1584.3 s (26.4 min). It holds 47,551 photometry samples per channel and 95,059 video frames, 47,530 of them on the 30-frames/s scoring grid. It contains 72 bouts: 16 approach, 19 investigation, 19 retrieval and 18 nest-based care. Sixteen of these form consecutive approach → investigation → retrieval → nest-based care sequences.

For those four behaviours the pads (*T*_pre/*T*_post) were 3/2.5, 2/1, 5/4 and 7/6.5 s. The before and after windows were each 2.5, 1, 4 and 6.5 s. The display limits were 5, 2, 6.5 and 16 s. All bouts entered every test. Each window equals the mean bout duration (2.28, 0.93, 3.60 and 6.33 s) rounded up to the next 0.5 s, exceeding the median (2.05, 0.87, 3.60 and 5.72 s) by 0.13–0.78 s. *T*_post equalled the after window, so every aligned trace covered it, and *T*_pre exceeded the before window by 0.5–1 s. Display limits include the longest bout of each behaviour (4.60, 1.57, 6.20 and 15.97 s), rounded up. The session was analysed with automatic test selection and without Holm correction.

The worked example singled out in **Figs 1, 2 and 4** is retrieval Trial 10 (frames 12481–12603; 4.07 s). The **Fig. 1** close-up around it covers 397.3–439.5 s. A script chose both by ranking the session’s complete behavioural sequences on display ΔF/F range. They are illustrative, not random. **Figs 1C and 1E** show the display ΔF/F; Figures 2–4 show the analysis ΔF/F. All panels are vector graphics generated by Node.js scripts that reproduce EthoTrace’s computations from the session files; re-running them reproduces their outputs byte for byte, and their values were checked against EthoTrace’s report and source. Outputs were cropped, relabelled and assembled in Adobe Illustrator; the legends identify redrawn interface panels and illustrations. These data document the workflow in one animal and are not presented as a biological finding.

The feature comparison in **Fig. 5** was scored from each tool’s primary publication, and for the behavioural event-logger additionally from its user guide. A feature not described in a source was scored as unreported rather than absent. Per-cell evidence and sources are given in Extended Data Table 1.

## Data and software availability

EthoTrace is distributed as a single HTML file that requires only a desktop browser, with no dependencies to install, no server component and no network requests. The tool, its documentation and the session data supporting this study are available at https://guangweizhang.com/tool-ethotrace.html, and from the corresponding authors on request.

## Author contributions

H.O. wrote the code and developed the tool, and performed the behavioural and fiber-photometry recording. C.T. and G.-W.Z. designed and supervised the project. C.T. and G.-W.Z. wrote the manuscript. All authors read and approved the final manuscript.

## Funding

This work was supported by the Whitehall Foundation (Grant 2026-08-034 to G.-W.Z.); the Brain & Behavior Research Foundation through a 2026 NARSAD Young Investigator Grant (Grant 34410 to C.T.); the VCU Convergence Team Planning Grant (to G.-W.Z.); and the Animal Models for the Social Dimensions of Health and Aging Research Network (NIH R24 AG065172; Pilot & Feasibility Award to G.-W.Z.).

## Competing interests

The authors declare no competing interests.

## Notes

https://guangweizhang.com/tool-ethotrace.html

### Competing Interest Statement

The authors have declared no competing interest.

